# SciViewer- An interactive browser for visualizing single cell datasets

**DOI:** 10.1101/2022.02.14.480435

**Authors:** Dhawal Jain, Sikander Hayat, Xinkai Li, Joydeep Charkaborty, Pooja Srinivasa, Michael H. Cho, Edwin K. Silverman, Hobert Moore, Rafael Kramann, Alexis Laux-Biehlmann

## Abstract

Single-cell sequencing improves our ability to understand biological systems at single-cell resolution and can be used to identify novel drug targets and optimal cell-types for target validation. However, tools that can interactively visualize and provide target-centric views of these large datasets are limited. We present SciViewer (**S**ingle-**c**ell **I**nteractive **View**er), a novel tool to interactively visualize, annotate and share single-cell datasets. SciViewer allows visualization of cluster, gene and pathway level information such as clustering annotation, differential expression, pathway enrichment, cell-type specificity, cellular composition, normalized gene expression and comparison across datasets. Further, we provide APIs for SciViewer to interact with publicly available pharmacogenomics databases for systematic evaluation of potential novel drug targets. We provide a module for non-programmatic upload of single-cell datasets. SciViewer will be a useful tool for data exploration and target discovery from single-cell datasets. It is available on GitHub (https://github.com/Dhawal-Jain/SciViewer).

## Introduction

Single-cell RNA-sequencing allows transcriptional profiling at single cell resolution. This technology has been instrumental in elucidating cellular heterogeneity, cell-type specific differential expression and transcriptional regulation programs, as well as rare cell types and cell states (Litviňuková *et al*., 2020; Tucker *et al*., 2020; Montoro *et al*., 2018) in different systems during homeostasis (Mayr *et al*., 2019), development (Raj *et al*., 2018; Srivatsan *et al*., 2021), disease (Kuppe *et al*., 2021, 2020; Parikh *et al*., 2019) and perturbed conditions (Srivatsan *et al*., 2020). More recently, single-cell sequencing has been useful in understanding COVID-19 infections and immune response (Melms *et al*., 2021). Subsequently technological advances in single-cell profiling have made it possible to study protein, chromatin assembly, surface receptors and perturbations at single cell resolution.

In translational research, single-cell sequencing has been used to understand changes in disease phenotype (Izar *et al*., 2020), discover mechanisms of treatment resistance (Bachireddy *et al*.; Chan *et al*., 2021; Jerby-Arnon *et al*., 2018) and drug tolerance (Aissa *et al*., 2021), and identify potential drug targets (He and Garmire, 2021). Additionally, single-cell sequencing can inform about cell-type specificity of a disease phenotype, reveal dysregulated molecular programs at single-gene and pathway levels, and identify relevant cell types for in vivo and in vitro target validation. Single-cell sequencing promises to herald a new era of high-throughput data-driven development or allocation of therapeutics. However, interpreting these vast amounts of sophisticated data requires interactive visualization tools that can enable non-programmatic access to the data for collaborative analyses across scientific disciplines.

Interactive visualization can be useful for quality control, cell-type annotation, and extraction of biologically relevant information from such datasets. As the number of studies, and cells and reads per study increase, visualization of millions of cells per study has become computationally challenging. Currently, a few solutions exist for single-cell sequencing visualization such as bbrowser (Le *et al*.), cellxgene (Megill *et al*., 2021), UCSF Cell Browser (UCSC Cell Browser) and the BROAD Single Cell Portal (https://singlecell.broadinstitute.org/single_cell). Importantly, most of these browsers have limited customization, and additional information about genes such as targetability, subcellular localization, and protein interaction networks cannot be incorporated. This leaves space for further development of interactive visualization portals. In this work, we present SciViewer, an open-source, customizable framework to interactively visualize single-cell datasets that can be used for target nomination and prioritization. From data upload to data analyses, SciViewer offers complete non-programmatic access to users. This will enable non-computational researchers to independently upload and analyze single-cell datasets. In SciViewer, a dataset can be visualized at multiple levels over major cell types, sub-types and compared to a second dataset. Additionally, multiple genes from the same datasets can be viewed and qualitatively compared. SciViewer is customizable, fast and can handle very large datasets. SciViewer is modular and in addition to visualizing single-cell datasets, it can be used to interact with other public resources. Furthermore, SciViewer functionality can be expanded to include single-cell modalities other than RNA-Seq. We envision that SciViewer will be a useful tool to prioritize genes for downstream experimental validation as potentially novel drug targets.

## 2. Materials and Methods

### Front-end

SciViewer is developed as an R-shiny (https://github.com/rstudio/shiny) application based on the Shiny framework developed by RStudio. It uses R-shiny and Javascript to organize the front-end of the application. SciViewer uses rasterly (https://github.com/plotly/rasterly), an open-source R library, for generating rasterized images of the 2D layout plots, which are then transferred to the client side for display. This significantly reduces the display time for client-side rendering. SciViewer front-end provides on-the-fly gene-based (one or multiple genes) visualization of single-cell transcriptomics data. Multiple 2D layout plots can also be generated to summarize gene-level expression, quality parameters, and gene-signatures. The application further allows interactivity with the images using custom javascript code, such that the users can hover over the images and get additional annotations on underlying cells.

SciViewer further summarizes the data using interactive bar charts and heatmaps. For enhanced interactive visualization, SciViewer uses open-source R-libraries such as plotly (https://github.com/plotly/) (Sievert, 2020), heatmaply (Galili *et al*., 2018) as well as a wrapper R-library highcharter (Kunst *et al*., 2017) for the licensed javascript module Highcharts (https://www.highcharts.com/). The license to use Highcharter should be organized by the end-users separately). Users can explore the data using metadata features available for the single cell study. In order to interact with public resources and perform functional enrichment of a set of genes, SciViewer uses API-based queries supported by gProfiler (Reimand *et al*., 2018). Additionally, using gage (Luo *et al*., 2009) and pathview (Luo and Brouwer, 2013) libraries, SciViewer provides functional enrichment of KEGG pathways and enhanced displaying of expression fold change information on the pathway maps.

### Back-end

SciViewer uses SQLite for storing single cell data. SciViewer provides a separate interactive shiny module for reading, digesting and creating SQLite databases from standard Seurat or h5ad data objects. Using this module, users can non-programmatically create a backend database for a single cell study and ingest it into SciViewer for interactive visualization. It is also possible to use the same module programmatically using the command line. For efficient storage and data retrieval, the column-oriented sparse data matrix (dgCMatrix) is aggregated into a 3-column data table (Supplementary Figure S1). Here, each table row represents a gene. The first and second columns represent aggregated column or row indices for a gene listing those cells where the gene has non-zero expression. The last column aggregates the normalized expression values. This format significantly reduces the space for storing the data, while allowing quick queries. Further, we have implemented functionality where the data can be stored remotely on cloud servers using MySQLite. In our experience, this allows separating the data from the application while still allowing quick real-time querying and display.

The input module of SciViewer allows calculation of cell-type markers and differentially expressed genes Briefly, SciViewer uses the ‘FindMarkers’ function from the Seurat package (Hao *et al*., 2021) to compute markers for cell-type or any other user-defined available study features such as disease state, clinical parameters, etc. Here, the cells are grouped based on selected attributes and the gene expression is compared against the remaining cells. Likewise, for calculating the differential gene expression, the data for each cell type is grouped between control and a user-selected test condition. Using the Wilcoxon test, the difference in expression is quantified and stored in the SQLite backend database.

While generating the SQLite database using SciViewer, the users can also precompute gene signatures for a selected set of genes. Currently, by default, the application allows calculating the expression signature for 186 KEGG (Kanehisa *et al*., 2017) and 662 WIKIPATHWAY (Slenter *et* al., 2018) gene sets as outlined in the C2 collection of MsigDB (version V7.4) (Subramanian *et* al., 2005).

## 3. Results

### A customizable visualization tool for single-cell multi-modal data analyses

We have developed SciViewer an open-access tool to interactively visualize single-cell data. Briefly, SciViewer is comprised of four streams: 1) data conversion and storage, 2) data visualization, 3) comparative analyses of single-cell datasets and 4) annotation of single-cell datasets using publicly available datasets to help aid in target prioritization. In addition to non-programmatic capabilities that can enable non-computational scientists to interactively explore datasets, SciViewer also provides APIs for programming access to its underlying resources.

### Input data conversion and storage

SciViewer uses an aggregated sparse matrix format to make data storage and retrieval more efficient for visualization purposes. SciViewer only stores information about genes that have a non-zero count for a given cell. Briefly, the sparse matrix (gene*cell format) is divided into two tables for a) storing metadata -where column index (column index identifies a unique cell) and other relevant information such as cell type and umap coordinates are stored for each cell barcode, and b) storing gene counts - where column index and count values are stored for each gene in the dataset (Supplementary Figure S1). SciViewer has a dedicated module for conversion of Scanpy or Seurat objects (Wolf *et al*., 2018; Hao *et al*., 2021) to an aggregated sparse matrix where users can upload their datasets by simply dragging and dropping their data objects, and provide information about metadata such as continuous and discrete variables, default 2D coordinate system to use, and cell-annotation columns in a non-programmatic manner (Figure 1). The input module then converts the dataset into an aggregated spare format and stores it as an SQLite database. This database is then used for interactive visualization by the SciViewer app.

**Figure 1:**
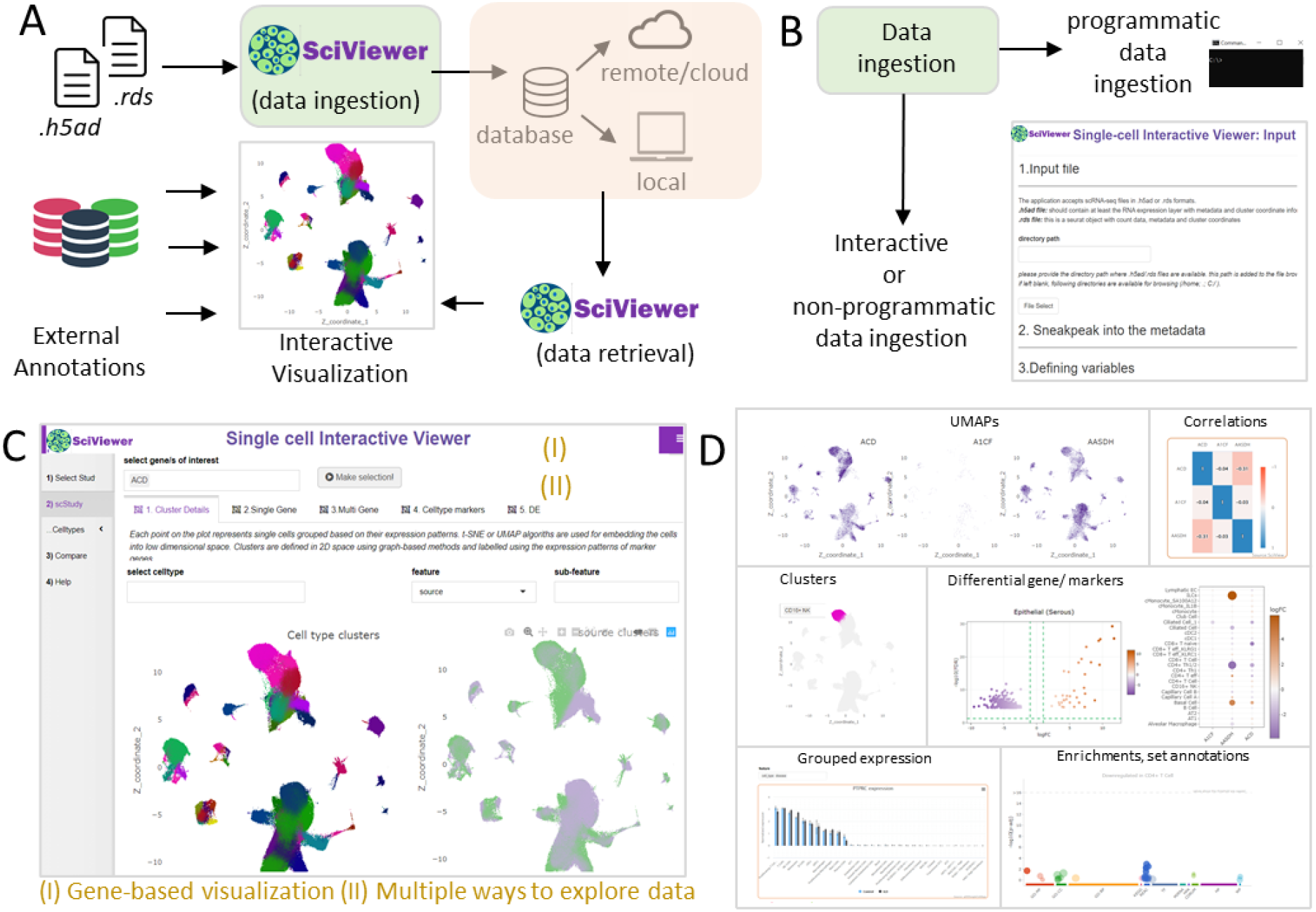
Single Cell Interactive Viewer (SciViewer) outline. (A) SciViewer can read Seurat-or H5AD-formatted single cell data to generate an underlying database. The browser also allows users to interactively visualize, explore and analyze the single cell data. (B) SciViewer allows both programmatic as well as an interactive platform for data upload and ingestion. (C) The front-end of the SciViewer browser allows gene-based visualization of the data. (D) Users can interact with the data in a range of different ways, such as plotting single or multi-gene expression, gene-expression correlations, cluster-based analyses, differential gene expression, expression grouped by covariates, and Gene-set enrichments.

### Visualization capabilities of SciViewer

SciViewer allows users to view single-cell data at different cell-type annotation levels. Separate tabs are provided to view different hierarchies of annotations (Figure 1). Cluster level view provides overall UMAP representation of different cell-types, which can be further refined to view either select or sub-types of major cell types. Gene-level view allows for overall 2D representation of the gene as UMAPs. Additionally, expression per cell type is shown as either bar or box-plots. Dotplots showing the percentage of cells expressing the gene of interest and its average expression are also available (Figure 1). Differentially expressed genes and pathway annotation for sorted gene lists can also be computed while ingesting the data in SciViewer or provided separately by the users. SciViewer also provides functions to compare two or more genes and visualize their expression as UMAPs, dotplots and correlation plots (Figure 1).

### Comparative analyses of single-cell datasets

Comparing different datasets can further enable analyses of single-cell datasets by providing insights into biological mechanisms under different conditions. For example, it can be useful to compare the cell-type specific expression of a given gene in a healthy reference with a disease dataset to understand compositional and expressional changes between the two conditions. Further, it can be helpful to confirm specificity of gene-expression of marker genes by comparing their expression across similar datasets. For target discovery, it can be highly relevant to compare the expression of potential novel targets in the appropriate cell-type in the tissue of interest to that of its expression in organs where dysregulation of that target might lead to toxicity and severe side-effects. SciViewer provides functionality to compare two datasets and view selected UMAPs, dotplots and barplots of genes across datasets (Figure 1).

## 4. Discussion

Multiple single-cell transcriptomics datasets are already in the public domain and with reducing costs and advancing technology, and many more datasets are becoming available (Svensson *et al*., 2020). These datasets cover different species and capture different aspects of biological diversity at the level of different individuals, anatomic sites and organs. Comparative analyses of these existing datasets can be useful for streamlining novel target and biomarker discoveries, and advancing translational research by providing robust results across different datasets capturing biologically diverse samples obtained from public sources. We have developed SciViewer as a customizable, fast, user-friendly, scalable and memory efficient tool to enable non-programmatic interactive visualization of single-cell transcriptomics datasets. Taken together, with back-end database storage, front-end visualization module, SciViewer enables finding, accessing and reusing single-cell datasets. SciViewer is compatible with both Seurat (Hao *et al*., 2021) and Scanpy (Wolf *et al*., 2018) data objects, can be easily set-up and used to visualize in-house data and compare it with existing publicly available datasets. In the future, external drug target annotation resources such as open targets (Koscielny *et al*., 2017), UniProt (UniProt Consortium, 2019), DisGeNET (Piñero *et al*., 2020), Omnipath (Türei *et al*., 2016) and other workflows such as Besca (Mädler *et al*., 2021) could be linked with SciViewer for target prioritization. SciViewer is modular and the codebase is provided in the public domain as open-access for users to further customize it to their specific needs. We expect that SciViewer will promote the FAIR principles (Clarke *et al*., 2019) of data sharing and improved interpretability of single-cell transcriptomics datasets.

## 5. Acknowledgements

We acknowledge the support of joint pulmonary drug discovery (J-PDD) and joint precision cardiology lab (J-PCL) of Bayer for - providing constructive feedback on this app. We further thank Dr. Craig Hersh, Dr. Vincent Carey, Dr. Dawn DeMeo, Dr. Pete Castaldi (CDNM, BWH, Harvard University), Dr. Edy Kim (BWH) and Dr. Florian Sohler, Dr. Justus Veerkamp (Bayer AG) and for valuable discussion and feedback on SciViewer.

---

**Supplemental Figure 1:**
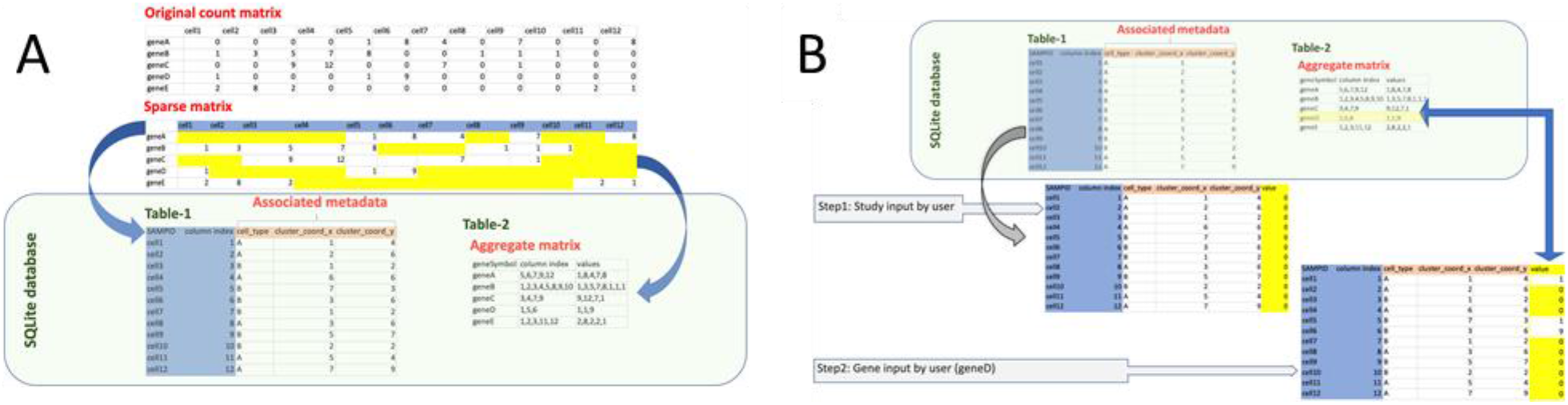
Data storage and retrieval by SciViewer. (A) Single cell count data are stored in the sparsematrix format inside Seurat or H5AD objects. SciViewer aggregates the counts into an aggregated matrix and a linked metadata file. The metadata file carries the cell information and additional annotations, while the aggregated matrix carries the column indices and the respective data values. (B) When a study is selected for browsing, SciViewer firsts retrieves the metadata file. Next, based on the queried gene, SciViewer retrieves relevant data from the aggregated matrix and attaches that to the metadata file. The metadata file is then used for further analyses, plotting 2D layouts and summarizing the data.

